# Multiple-object tracking as a tool for parametrically modulating memory reactivation

**DOI:** 10.1101/109256

**Authors:** J. Poppenk, K.A. Norman

## Abstract

Converging evidence supports the “non-monotonic plasticity” hypothesis that although complete retrieval may strengthen memories, partial retrieval weakens them. Yet, the classic experimental paradigms used to study effects of partial retrieval are not ideally suited to doing so, because they lack the parametric control needed to ensure that the memory is activated to the appropriate degree (i.e., that there is some retrieval, but not enough to cause memory strengthening). Here we present a novel procedure designed to accommodate this need. After participants learned a list of word-scene associates, they completed a cued mental visualization task that was combined with a multiple-object tracking (MOT) procedure, which we selected for its ability to interfere with mental visualization in a parametrically adjustable way (by varying the number of MOT targets). We also used fMRI data to successfully train an “associative recall” classifier for use in this task: this classifier revealed greater memory reactivation during trials in which associative memories were cued while participants tracked one, rather than five MOT targets. However, the classifier was insensitive to task difficulty when recall was not taking place, suggesting it had indeed tracked memory reactivation rather than task difficulty per se. Consistent with the classifier findings, participants’ introspective ratings of visualization vividness were modulated by MOT task difficulty. In addition, we observed reduced classifier output and slowing of responses in a post-reactivation memory test, consistent with the hypothesis that partial reactivation, induced by MOT, weakened memory. These results serve as a "proof of concept” that MOT can be used to parametrically modulate memory retrieval – a property that may prove useful in future investigation of partial retrieval effects, e.g., in closed-loop experiments.

## Introduction

Although retrieval from episodic memory is thought to be obligatory and complete (Moscovitch, Cabeza, Winocur, & Nadel, 2016), control processes may operate on the product of retrieval to induce states of partial memory reactivation. According to the non-monotonic plasticity hypothesis (Newman & Norman, 2010), such partial memory reactivations can weaken memory representations, even though full reactivations can strengthen them. Non-monotonic learning is supported by various lines of evidence: for example, the large and growing cognitive literature on retrieval-induced forgetting (Murayama, Miyatsu, Buchli, & Storm, 2014); neurophysiological evidence of moderate, but not high, levels of depolarization leading to weakening (Artola, Brocher, & Singer, 1990; Hansel, Artola, & Singer, 1996); neural models of synaptic plasticity (Norman, Newman, Detre, & Polyn, 2006); and impaired subsequent memory for events shown to be partially activated by EEG and fMRI (e.g., Detre, Natarajan, Gershman, & Norman, 2013; Kim, Lewis-Peacock, Norman, & Turk-Browne, 2014; Lewis-Peacock & Norman, 2014; Newman & Norman, 2010; Poppenk & Norman, 2014; Wimber, Alink, Charest, Kriegeskorte, & Anderson, 2015). But as empirical evidence for non-monotonic learning accumulates, what tools are needed to further advance the field?

A key limitation of existing studies that have been used to characterize non-monotonic learning is that they rely on *naturally-occurring variability* within experimental conditions. For example, Detre et al. (2013) used a think / no-think paradigm (Anderson & Green, 2001), measured (on each trial) how much participants thought of “no think” memories that they were not supposed to be retrieving, and related this within-condition variance to subsequent memory. In that study, the naturally-occurring distribution of memory activation values was wide enough to characterize the full U-shaped curve (i.e., no memory change for very low activation, memory weakening for moderate activation, and memory strengthening for higher levels of activation). Crucially, there is no guarantee that any given study will obtain broad enough “coverage” of the range of memory activation values to trace out the full curve (indeed, in Detre et al., 2013, there were substantially more activation values towards the middle of the activation range than towards the high and low extremes; we were lucky that there were enough observations to run the analysis). Existing paradigms (e.g., think / no-think) tend to use binary manipulations of memory activation, which further limits the range of activation values sampled in the experiment. What if we needed to obtain partial memory reactivation occurring halfway between that induced by "think" vs. "no-think" instructions? It would be a great benefit to have a finer-grained “dial” that we could adjust in experiments to increase the range of memory activation values that we sample. This capability could, for example, allow therapists treating patients with post-traumatic stress disorder to more effectively reactivate memories to levels known to induce memory weakening.

In a recent study (Poppenk & Norman, 2014), we set out to parametrically modulate memory activation using an adaptation of an rapid serial visual presentation (RSVP) design that we called "The Great Fruit Harvest”. Participants associated word memory cues with pictures of bedrooms; these word memory cues were then embedded in an RSVP stream that participants were monitoring for fruit words (note that none of the word memory cues were themselves fruit words). To manipulate the degree of memory reactivation, we varied how long the word cues were presented in the RSVP stream. Reactivation of associated scene memories in response to these cues was tracked using an fMRI pattern classifier trained to detect scene information. The cue-duration manipulation was successful in generating differential memory effects: Compared to longer (2000 ms) word-cue presentations, brief (600 ms) word-cue presentations led to lower levels of memory activation and more memory weakening. In light of these results, we think that the word-cue duration manipulation has promise. However, in this paradigm, recall elicited by a memory cue is always task-irrelevant, as it distracts from looking for fruit words. Thus, associated cues should always be suppressed, potentially making it difficult to trace out the full U-shaped curve.

Here, we present an alternative approach to generating parametrically scalable memory reactivation. This approach is based on the idea that it is critical to make memory retrieval an *explicit part of the task*, such that participants will not automatically suppress strong memory retrieval. Also, instead of varying the strength of the memory cue (as in our RSVP design), we varied the cognitive demands of a distractor task that competed with memory retrieval. The distractor task we selected was multiple object tracking (MOT; Pylyshyn & Storm, 1988). Briefly, participants were required to track a variable number of MOT targets within a moving dot field over an eighteen-second interval, with dots moving at a speed calibrated to each participant’s visuospatial ability. Concurrently with this task, participants were asked to visualize the scene associate of a word cue presented in the centre of the screen, and to provide ongoing ratings concerning the integrity of their mental visualization. Throughout instruction and practice for this task, we emphasized that MOT dot-tracking should take precedence, and that visualization should only be "squeezed in" using available mental resources. To further emphasize this point, we provided feedback on dot-tracking accuracy after every trial. We selected this combination of tasks because, as a visuospatial task, we anticipated that MOT would compete for the visual resources required for visualization of mental imagery (Phillips & Christie, 1977). Furthermore, a key property of MOT is that participants need to *continuously* attend to the task – any lapse of attention will break the train of observations linking each dot to its targetness, making it impossible to solve the correspondence problem as required for successful responding (Pylyshyn, 2004). Accordingly, we reasoned that the MOT task would both a) impair visualization of any retrieved information; and b) make it difficult for participants to momentarily switch out of the MOT task to apply full concentration to visualization.

We predicted that, by varying the number of MOT targets participants were required to monitor, we would parametrically modulate resources available for mental visualization, and would observe corresponding variation in memory reactivation. We further predicted that partial memory activation induced by this procedure would lead to forgetting effects consistent with the non-monotonic plasticity hypothesis and its supporting literature.

## Methods

### Overview

The experiment contained several main phases (see Table 1 and Fig. 1): MOT difficulty calibration (Phase 1), paired-associate training (Phase 2), memory reactivation (Phase 5), and pre-and post-reactivation memory tests (Phases 4 and 6). In addition, a functional localizer was collected to assist with pattern classification analysis (Phase 3). This design was modeled after that used by Poppenk and Norman (2014), but it incorporated a novel method for reactivating memories (Phase 5), as well as a novel procedure for training a classifier sensitive to memory reactivation (Phase 3). We employed an MOT task in which participants tracked moving MOT target dots among a set of identically-colored moving lure dots (Pylyshyn & Storm, 1988) while centrally fixating on a verbal memory cue. We attempted to modulate memory reactivation by altering the number of MOT target dots that participants were required to track in the MOT task. We also attempted to train a classifier that could be used to provide additional insight into memory reactivation. Our hypotheses concerned the effectiveness of our protocol at modulating memory reactivation (Phase 5), the ability of our classifier to measure this modulation (Phase 3), and whether partial reactivation as measured by our instruments would successfully induce forgetting as observed in a post-reactivation memory test (Phase 6), as partial reactivation in other paradigms has been shown to do.

**Table 1.**
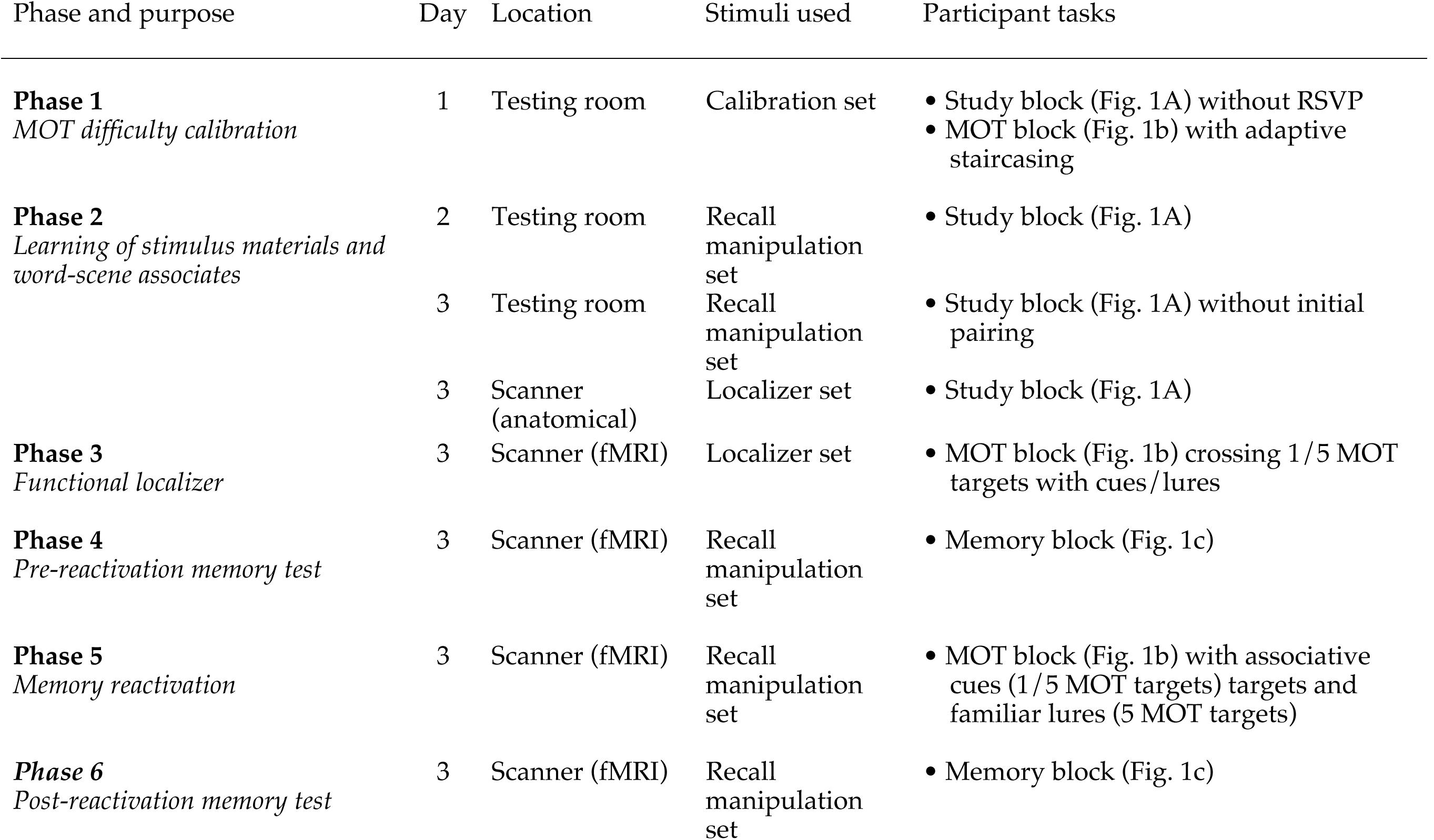
Schematic of main experimental phases.

| Phase and purpose | Day | Location | Stimuli used | Participant tasks |
| --- | --- | --- | --- | --- |
| **Phase 1**<br>*MOT difficulty calibration* | 1 | Testing room | Calibration set | • Study block (Fig. 1A) without RSVP<br>• MOT block (Fig. 1b) with adaptive staircasing |
| **Phase 2**<br>*Learning of stimulus materials and word-scene associates* | 2 | Testing room | Recall manipulation set | • Study block (Fig. 1A) |
|  | 3 | Testing room | Recall manipulation set | • Study block (Fig. 1A) without initial pairing |
|  | 3 | Scanner (anatomical) | Localizer set | • Study block (Fig. 1A) |
| **Phase 3**<br>*Functional localizer* | 3 | Scanner (fMRI) | Localizer set | • MOT block (Fig. 1b) crossing 1/5 MOT targets with cues/lures |
| **Phase 4**<br>*Pre-reactivation memory test* | 3 | Scanner (fMRI) | Recall manipulation set | • Memory block (Fig. 1c) |
| **Phase 5**<br>*Memory reactivation* | 3 | Scanner (fMRI) | Recall manipulation set | • MOT block (Fig. 1b) with associative cues (1/5 MOT targets) targets and familiar lures (5 MOT targets) |
| **Phase 6**<br>*Post-reactivation memory test* | 3 | Scanner (fMRI) | Recall manipulation set | • Memory block (Fig. 1c) |

**Fig. 1.**
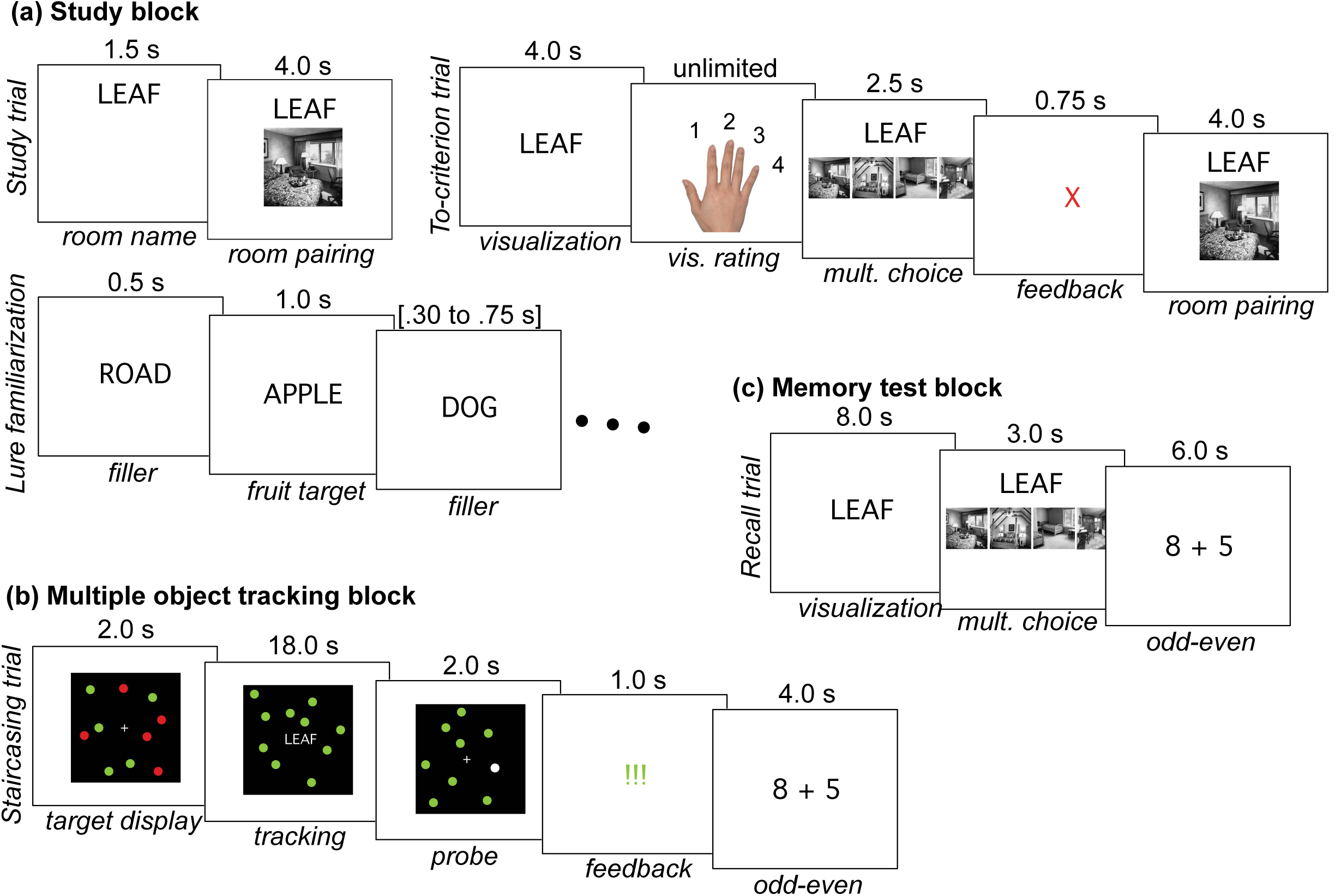
Trial types for Phases described in Table 1. In study blocks (a), participants first studied word-scene associate pairs by viewing them once. Then, they learned the pairs to criterion: upon being presented with a cue word, they rated the amount of detail in their mental visualization of the word, completed a 4AFC decision for its scene associate, and were presented with feedback (incorrect items were repeated). Finally, participants were familiarized with words that had no associates. Participants monitored a stream of words for embedded fruit items, and pushed a button when a fruit item appeared. Filler words were used as lures in later memory tests. In later memory test blocks, words from the word-scene associations were used to cue mental visualization. Filler words from the RSVP task were used during familiarization as familiar lures. In MOT blocks (b), participants completed cued visualization while simultaneously tracking target dots in an MOT task. Each time a central fixation dot turned red, participants reported current levels of visualization. After the trial, participants were given feedback and completed a series of odd-even judgments. In memory test blocks (c), participants completed cued visualization of studied words’ associates as well as multiple choice for their associates.

### Participants

24 right-handed volunteers participated in the experiment (16 female, age *M* = 20.5 years, *SD* = 1.6 years). Six additional participants were unable to retain the positions of five nearly immobile MOT targets during Phase 1, and were not invited to complete the experiment (details below). One participant was excluded because of excessive in-scanner motion, leaving a total of 23 participants. All were native English-speakers between 18 and 25 years of age with normal or corrected-to-normal vision and hearing. Participants were screened for neurological and psychological conditions, and received financial remuneration. The protocol was approved by the Institutional Review Board for Human Subjects at Princeton University.

### Stimuli

Participants learned three sets of word-scene pairings. A *calibration set* of eight pairs was used during the MOT difficulty calibration phase (Phase 1); a *localizer set* of sixteen pairs was used in the functional localizer (Phase 3); and a *recall manipulation set* of 30 pairs was used for testing experimental hypotheses (Phases 2, 4, 5 and 6; see Table 1). Words were concrete, imageable nouns randomly sampled from a pool of 7000 nouns drawn from the MRC Psycholinguistic Database (Coltheart, 1981; mean length = 6.3 letters; mean concreteness = 571.5; mean imageability = 561.3; mean Thorndike-Lorge verbal frequency = 241.68). Words were filtered to exclude nouns semantically related to rooms. Paired scenes were grayscale bedroom interiors drawn from Detre et al. (2013). Each participant received a different random pairing of words and images.

There were also two sets of words used as lures: a set of 16 used during the functional localizer phase (Phase 3), and a set of five used during the MOT phase (Phase 5). These words were randomly sampled from the same pool as above.

All text in the experiment was presented in black Geneva font (height = 0.8° visual angle) on a white background. All images in the experiment were the same size (9.0° × 9.0° visual angle) and normalized with respect to their luminance using the procedure described in Detre et al. (2013).

### Procedure

#### Phase 1. MOT difficulty calibration

Because visuospatial ability varies considerably across individuals, yet we desired to modulate MOT in a way that would be comparably distracting for each participant, it was necessary to calibrate the difficulty of MOT to the saturation point of each participant’s abilities. We did so by beginning the experiment with a phase in which we used a staircasing procedure to adjust the speed at which the MOT task ran. In this phase, which took place in a behavioral testing room at least one day prior to the rest of the experiment, participants first studied the *calibration set* of eight word-scene associates (Table 1; Fig. 1a). After studying these pairs to criterion, they began the MOT staircasing task.

To study the word-scene associates, the eight pairs were presented once. Then, the order was randomized, and the eight pairs were presented again (Fig. 1a). Participants were told that a memory test would follow and that, to make stronger memories, they should treat the cue word paired with each bedroom image as the name of that “hotel room”. They were told they should imagine the most creative, distinctive possible explanation for how each “hotel room” got its name. Cue words were presented for 5.5 s; 1.5 s after each cue word onset, the scene image also appeared below the word. A fixation period 0.75 s in duration separated trials.

Next, participants completed a train-to-criterion memory test (Fig. 1a). Each trial incorporated three parts. First, a cue word was presented for four seconds, during which time participants were instructed to visualize the associated scene in as much detail as possible. Next, they were asked to rate their visualization on the following scale: 1: no room-related imagery or a generic room with no distinguishing features; 2: room with a specific distinguishing feature; 3: room with multiple specific distinguishing features; 4: complete image. After a subjective response was registered, the associated scene image plus scenes randomly selected from three other studied pairings were presented in random order from left to right. Participants had three seconds to select the scene associated with the cue word via a button press. If a correct response was entered before the deadline, green exclamation points were presented for 0.75 s. Otherwise, a red “X” was presented for 0.75 s, followed by presentation of the cue word with the correct scene image for four seconds. A five second fixation cross separated each trial. Each item remained in the list until it received a correct multiple-choice response, at which point it was dropped from the study set. The order of the (remaining) pairs in the list was randomly shuffled after each pass through the list.

In the final section of the calibration session, participants completed eight practice MOT trials to become familiar with the task, then 72 additional MOT trials in which we adjusted the speed of the task based on their ability (Fig. 1b). MOT trials consisted of a black central square (20° x 20° visual angle) containing ten dots (each 1.5° diameter). In each trial, each of the dots was assigned a random, non-overlapping starting position in the square, and five of the dots were shown in red (“targets”), while the remaining dots were shown in green (“non-targets”). In addition, a fixation cross was shown in white. After a two-second exposure duration, all dots were presented in green and began moving (their movement pattern was complex and is described in detail below). Participants were asked to mentally track which dots were originally the red target dots for eighteen seconds. In addition, the central fixation cross was replaced by a cue word from the *calibration set* in a white font. Participants were asked to visualize the associated room in as much detail as possible. Every four seconds, the cue word was switched to a red-colored font to signal that participants should make a visualization rating (of the same type performed in the memory test). After a response was detected or two seconds – whichever came first – the cue word was switched back to a white-colored font. At the end of the trial, all dots froze and one dot was presented in white. Participants were asked to indicate with a button-press whether this “probe” dot was originally a target or a non-target. After three seconds elapsed, participants were given feedback for one second indicating whether they were correct or incorrect (using the same format as in the memory test). Participants were also asked to always prioritize the MOT task over the visualization task, "squeezing in" visualization only when it would not compromise dot-tracking. We explained that while we were interested in their visualization, it came second to dot-tracking, as incorrect dot-tracking trials would have to be discarded. Finally, to disrupt any post-trial visualization of the cued scene imagery, participants completed two trials of an odd-even task: two digits were presented together with an addition symbol between them for 1.9 s, and participants were to indicate with a button-press whether the sum of the numbers was odd or even. This text was presented in white, but when a correct response was detected, the font was switched to green; when an incorrect response was detected, the font was switched to red. A total of 0.1 s of central fixation followed each odd-even question, and after both odd-even trials, four seconds of central fixation preceded the next MOT trial. Together, these elements comprised a total duration of 32 s per MOT trial.

In the MOT task, each dot began with independent *x-* and *y*-dimension starting velocities consisting of values sampled from a continuous random distribution (*x, y* = [-.5 to .5]) and multiplied by a velocity *v,* measured in visual degrees per second. After each frame, dot motion was recomputed. In particular, random *x* and *y* values were again sampled from a continuous random distribution (-.5 to .5), scaled by *v*, and added to each dot’s velocity. An additional vector was added to each dot’s velocity based on its position relative to that of other dots and the center of the square: all dots generated a “repulsion field” to reduce collisions with other dots. The repulsion effect of a dot on all other dots was calculated as 0.1*v* over the squared distance between them (yielding exponentially larger repulsion values as dots grew closer). A speed limit was enforced by capping dot velocity at an absolute velocity of 2*v* on each dimension. Finally, when a dot collided with another dot or the perimeter of the square (with collisions defined as occurring 1.25 diameters away from the center of a dot), *v* on the dimension in which the collision occurred was multiplied by −1 (yielding a “reflection” on that dimension).

We adjusted the parameter *v* throughout Phase 1 while presenting new frames at a rate of 30 per second. After participants completed eight practice trials at an initialization speed (1.0°/s), we adjusted *v* depending on whether participants succeeded in that trial, using the Quest adapted staircasing algorithm (Watson & Pelli, 1983) to calculate the optimal adjustments to identify the speed threshold at which participants would succeed 85% of the time (beta=3.0, delta=0.1, gamma=0.5, grain=0.2°/s, range=5.0°/s). This speed was used for all subsequent MOT-based tasks completed by the participant for the remainder of the experiment. As is typical in MOT experiments, only participants whose threshold fell above a given minimum (in our case, 0.5°/s) were invited to continue.

After completing the MOT task but before going home, participants completed a short study session and memory test for 40 proverbs. This memory test was conducted to test hypotheses that were unrelated to the current study, and because the test came after all other tasks on the difficulty calibration day, corresponding methodological details are not reported here.

#### Phase 2. Learning of stimulus materials and word-scene associates

The MOT difficulty calibration session was conducted well ahead of the rest of the experiment to ensure, prior to scanner scheduling, that participants would qualify for further experimentation (Phase 1 to 2 latency *M* = 8.41 days; *SD* = 9.65 days; range = 1-27 days). Participants studied the *recall manipulation* set of 30 paired-associates, then performed a train-to-criterion memory task with those pairs. Initial study and the train-to-criterion test were conducted in the same manner as in Phase 1 (except that the 30-item *recall manipulation set* was employed; Table 1). Participants quickly learned the 30 paired associates to criterion levels (*M* = 37.0 trials, *SD* = 7.5 trials). By later reactivating scene associates of the word cues within this set by differing amounts (in Phase 5, during an MOT task), we would attempt to weaken these memories. All of Phase 2 took place in a behavioral testing room, away from the scanner.

After the train-to-criterion task, participants were given a 60 s RSVP task in which they viewed fruit words and non-fruit words while responding to fruit words with a button press. Seven non-fruit words were presented repeatedly in random order, with the duration of each presentation sampled from a uniform distribution with limits of 0.30-0.75 s. Three fruit words were also presented for one second during the task, appearing at random intervals but no sooner than eight seconds after a previous fruit target. Participants were given feedback on their performance at the end of the task. The purpose of exposing participants to the non-fruit words in the RSVP task was to familiarize these words (without linking them to a scene associate) so that five of them could be used as familiar lures in Phase 5 of the experiment (and two as practice items). Although each of the word presentations during RSVP was brief, the cumulative presentation time of each familiar lure across all RSVP presentations (M=6.9 s) was matched to the total presentation time of cue words during study trials (7.0 s).

#### Phase 3: Functional localizer

The goal of this phase was to obtain a clean neural signal associated with cued retrieval of scenes that was insensitive to changes in MOT task difficulty. Approximately one day after learning the materials comprising the recall manipulation set (Phase 2 to 3 latency *M* = 23.2 hours; *SD* = 3.6 hours; range = 16.1-30.2 hours), participants returned for a third session. This entire session took place inside an fMRI scanner and began with a practice version of the localizer scan (see procedure description below), using paired associates from the *difficulty calibration set* learned in Phase 1. Then, while a high-resolution anatomical scan was collected, participants studied the 16-item *localizer* set of paired-associates (Table 1), completed a train-to-criterion memory task for those materials, and completed an RSVP task (Fig. 1a). This served to provide participants with a newly-acquired set of paired-associate memories and familiar lure words for use with a functional localizer. Tasks were presented in the same manner as in Phase 2 Here, the RSVP task involved showing sixteen (non-fruit) words plus eight fruit targets; the sixteen non-fruit words later served as familiar lures in the localizer. The RSVP task lasted 136 s and required participants to make button-presses on an MR-compatible keyboard.

The localizer consisted of a 32-item MOT task similar to that in Phase 1 (Fig. 1b). However, the centrally presented cues in the task consisted of the 16 localizer-set paired-associate cues (“cues”) and the 16 localizer-set familiar words without scene associates from the RSVP task (“familiar lures”). Each of these stimuli was presented once and their order was randomized. Half of the cue trials and half of the lure trials involved tracking only one MOT target, whereas the remaining trials involved tracking of five MOT targets. In this way, we crossed the presence of memory signal (cues vs. familiar lures) with task difficulty (1 vs. 5 MOT targets). This allowed us to use the associated fMRI data for training a classifier to identify memory signal (associative recall) in a manner that generalized across task difficulty level (number or MOT targets). This task took 17.2 minutes (517 fMRI volumes) to complete.

#### Phases 4 and 6: Pre- and post-MOT memory tests

In Phases 4 and 6, test items were all 30 cues from the recall modulation set (Table 1). No familiar lure words were required in these phases, as a four-alternative forced choice (4AFC) task (in which foils were the associates of other cues) was used to obtain an objective measure of memory performance (Fig. 1c). On each test trial, participants were first presented with a memory cue for eight seconds, during which they were instructed to visualize the associated scene in as much detail as possible. Next, the multiple-choice prompt was presented, along with four scenes (as in Phase 1). Participants had 3 s to choose, using a button press, which scene went with the cue word. This period was followed by three odd-even questions; as in Phase 1, these lasted 1.9 s each with a preceding fixation interval of 0.1 s. Questions were followed by five seconds of central fixation. No feedback was presented, and the pace of the experiment did not vary based on participant responses. The full group of 30 trials took 11.2 minutes (335 fMRI volumes) to complete. We analyzed accuracy and reaction time for all trials of each memory test.

#### Phase 5: Controlled memory reactivation in an MOT task

The goal of Phase 5 was to repeatedly elicit controlled levels of memory reactivation by placing word cues in MOT trials that featured different levels of difficulty. As in the Phase 3 localizer, difficulty was manipulated by requiring participants to track either 1 or 5 MOT targets. Of the 30 word-scene pairs in the recall manipulation set, ten pairs were assigned to the *associative cue* (*1 MOT target*) condition, which was intended to elicit the strongest reactivation; ten pairs were assigned *associative cue* (*5 MOT targets*) condition, which was designed to elicit weaker reactivation due to increased distraction from the MOT task; and ten pairs were *omitted* from this phase, so that they would not undergo any reactivation. Each fMRI run included one MOT trial for each of: 1) the ten cues from the associative cue (1 MOT target) condition; 2) the ten cues from the associative cue (5 MOT targets) condition; and 3) five familiar words from the Phase 2 RSVP task (familiar lure condition). The sequence of these trials was randomized for each fMRI run. Three runs were completed, each lasting 13.5 min (405 fMRI volumes).

### fMRI data collection

Scanning was performed using a 3 Tesla whole-body Skyra MRI system (Siemens, Erlangen, Germany) at Princeton University in Princeton, New Jersey. T1-weighted high-resolution MRI volumes were collected using a 3D MPRAGE pulse sequence optimized for gray-white matter segmentation, with slices collected in the AC-PC plane (176 sagittal slices; 1 mm thick; FOV = 256 mm; 256 × 256 matrix; TR = 2530 ms; TE = 3.37 ms; flip angle = 9°). All functional MRI scans were collected using T2*-weighted echo-planar image (EPI) acquisition (34 axial oblique slices; 3 mm thick; FOV = 192 mm; 64 × 64 matrix; TR = 2000 ms; TE = 33.0 ms; flip angle = 71°; 2x IPAT acquisition). A T1 FLASH and fieldmap image were also collected using these parameters to assist with coregistration of fMRI volumes to brain anatomy, and to correct spatial distortions.

### fMRI pre-processing

For each functional image, we computed the linear transformation required to coregister the image to the mean image of the first functional run, yielding an affine motion correction matrix. Using a fieldmap image, we also computed the warp field necessary for correction for spatial distortion of functional images, then combined the two transformations and applied them to the functional data in a single spatial transformation step. Then, we applied a high-pass filter (full-width half maximum = 160 s) and de-spiking algorithm to each voxel (3dDespike, AFNI).

We next segmented anatomical images to obtain participant-specific functional masks. We performed this segmentation in a semi-automated fashion using the FreeSurfer image analysis suite, which is documented and available online (v5.1; http://surfer.nmr.mgh.harvard.edu) with details described elsewhere (e.g., Fischl et al., 2004). Briefly, this processing includes removal of non-brain tissue using a hybrid watershed/surface deformation procedure, automated Talairach transformation, intensity normalization, tessellation of the gray matter white matter boundary, automated topology correction and surface deformation following intensity gradients, parcellation of cortex into units based on gyral and sulcal structure, and creation of a variety of surface based data including maps of curvature and sulcal depth. Manual quality control checks were performed. We resampled FreeSurfer segmentations to functional image space for use as anatomical masks. Based on meta-analysis implicating precuneus, fusiform, parahippocampal, inferior frontal, cingulate, inferior parietal, and superior parietal gyri in episodic memory recall (Spaniol et al., 2009), we assembled these segmentations into a “recall” mask for use with subsequent analyses.

### Classifier training

To support our analyses linking memory reactivation to later memory outcomes, we aimed to establish an ongoing, incidental measure of memory reactivation. In pilot testing, using data from a functional localizer phase, we attempted to train a classifier sensitive to multiple visual categories (faces, scenes, cars, and words; Spiridon & Kanwisher, 2002). We hoped to use the classifier to measure reactivation of scene unit in response to word cues that participants had previously studied in conjunction with scenes. We have used this indirect approach of monitoring memory reactivation previously (e.g., Detre et al., 2013; Poppenk & Norman, 2014) and it has become relatively common in the literature. However, we found that our MOT task, with multiple moving dots, would consistently and inappropriately elicit activity in the scene unit, perhaps because the composite of multiple independent objects within a black frame constituted a "scene" in a neural framework. This bias was apparent even when the classifier was trained with the MOT task active and the visual categories presented as a backdrop, and was sufficiently prominent as to prevent us from measuring memory reactivation in the typical manner.

To sidestep this issue, we adopted a classifier training protocol focused on the presence of an associative recall signal, similar to that developed by Rissman, Greely, and Wagner (2010). Rather than attempting to measure neural evidence for activation of scenes in the brain (i.e., memory content) we instead searched for neural evidence of episodic memory retrieval (i.e., memory operations). In particular, we trained a classifier to distinguish MOT trials incorporating words that were cues for previously-studied scene associates on the one hand (the "cue" condition), against words that were merely familiar due to prior exposure on the other (the "familiar lure" condition; it is worth noting that our designation of trials as "cues" or "familiar lures" was based on the experimental treatment of the word, rather than the subjective experience of the participant). Importantly, we incorporated equal numbers of 5- and 1- MOT target trials in each of the two memory conditions (associative cue and familiar lure). By including this MOT-difficulty manipulation but making it irrelevant (orthogonal) to the distinction being learned by the classifier (associative cue vs. familiar lure), we hoped to encourage the classifier to focus on recall-related variance and to ignore variance directly related to the number of MOT targets. This is a tricky issue: The point of having participants do the MOT task simultaneously with the memory task is to affect the level of memory activation, and we want the classifier to be sensitive to these *indirect* effects of MOT on recall. At the same time, we definitely did not want the classifier to be *directly* sensitive to the features of MOT, which is why we included a MOT-difficulty manipulation in our classifier training regime. The procedure that we chose can be viewed as conservative: by training the classifier to be insensitive to features of the MOT task, we ran the risk of making the classifier insensitive to indirect effects of MOT on recall, with the benefit that – if they are obtained – we can more clearly interpret these effects as pertaining to variance in recall (as opposed to variance in the surface features of the MOT task). Below (in the *Results*), we present several key analyses showing that the classifier has the properties that we sought. In Phase 3, we found that classifier output on familiar lure trials was *not* sensitive to the number of MOT targets (showing that, on trials where associative recall was *not* taking place, the classifier was not affected by properties of the MOT task); and in Phase 5, we found that classifier output on associative-cue trials *was* sensitive to the number of MOT targets (showing that, when recall *was* taking place, it was modulated in the anticipated fashion by the MOT task).

We performed our classifier analysis in Matlab using functions from the Princeton Multi-Voxel Pattern Analysis (MVPA) Toolbox (Detre et al., 2006; available for download at http://www.pni.princeton.edu/mvpa/), in the same manner as described in Poppenk & Norman (2014; see also Norman, Polyn, Detre, & Haxby, 2006, for a discussion of the logic and affordances of MVPA). Classifier training was performed separately for each participant using a ridge regression algorithm, which is sensitive to graded signal information (such as might be associated with intermediate states of memory reactivation). Ridge regression learns a ß weight for each input feature (voxel) and uses the weighted sum of voxel activation values to predict outcomes (in this case, a binary vector indicating which task is associated with each volume). The ridge regression algorithm optimizes each ß to simultaneously minimize both the sum of the squared prediction error across the training set and also the sum of the squared ß weights (technical details are described elsewhere; see Hastie, Tibshirani, & Friedman, 2001, and Hoerl & Kennard, 1970). A regularization parameter (λ) determines how strongly the classifier is biased towards solutions with a low sum of squared ß weights; when this parameter is set to zero, ridge regression becomes identical to multiple linear regression. The solution found by the classifier corresponded to a ß map for each regressor describing the spatial pattern that best distinguished that regressor’s condition from other conditions (with regularization applied).

We provided as input to the classifier all of the grey-matter voxels that fell within the "recall" mask described above, and set our ridge regression penalty parameter (λ) to 1. We also input a training regressor describing the presentation of cue words and familiar lure words, shifting our regressor by four seconds (i.e., two TRs) to accommodate hemodynamic lag effects associated with the blood-oxygen level dependent response in fMRI data.

To evaluate the effectiveness of this classifier at distinguishing between categories of images based on patterns of activity within the recall mask, we held out portions of the data when training for classifier testing (Kriegeskorte, Simmons, Bellgowan & Baker, 2009). The localizer was divided into eight “folds”, each of which contained one of the four trial types (associative cues with 5 MOT targets, associative cues with one MOT target, familiar lures with 5 MOT targets, and familiar lures with 1 MOT target). We left out one fold of each type (i.e., one eighth of the examples) on each iteration. As a reminder, although there were four types of trials, we trained on only two categories (cue and familiar lure trials), collapsing across number of MOT targets. Collapsing across folds, mean classifier accuracy was above chance across participants (Fig. 2b), BSR = 3.11, *P* < 0.005.

**Fig. 2.**
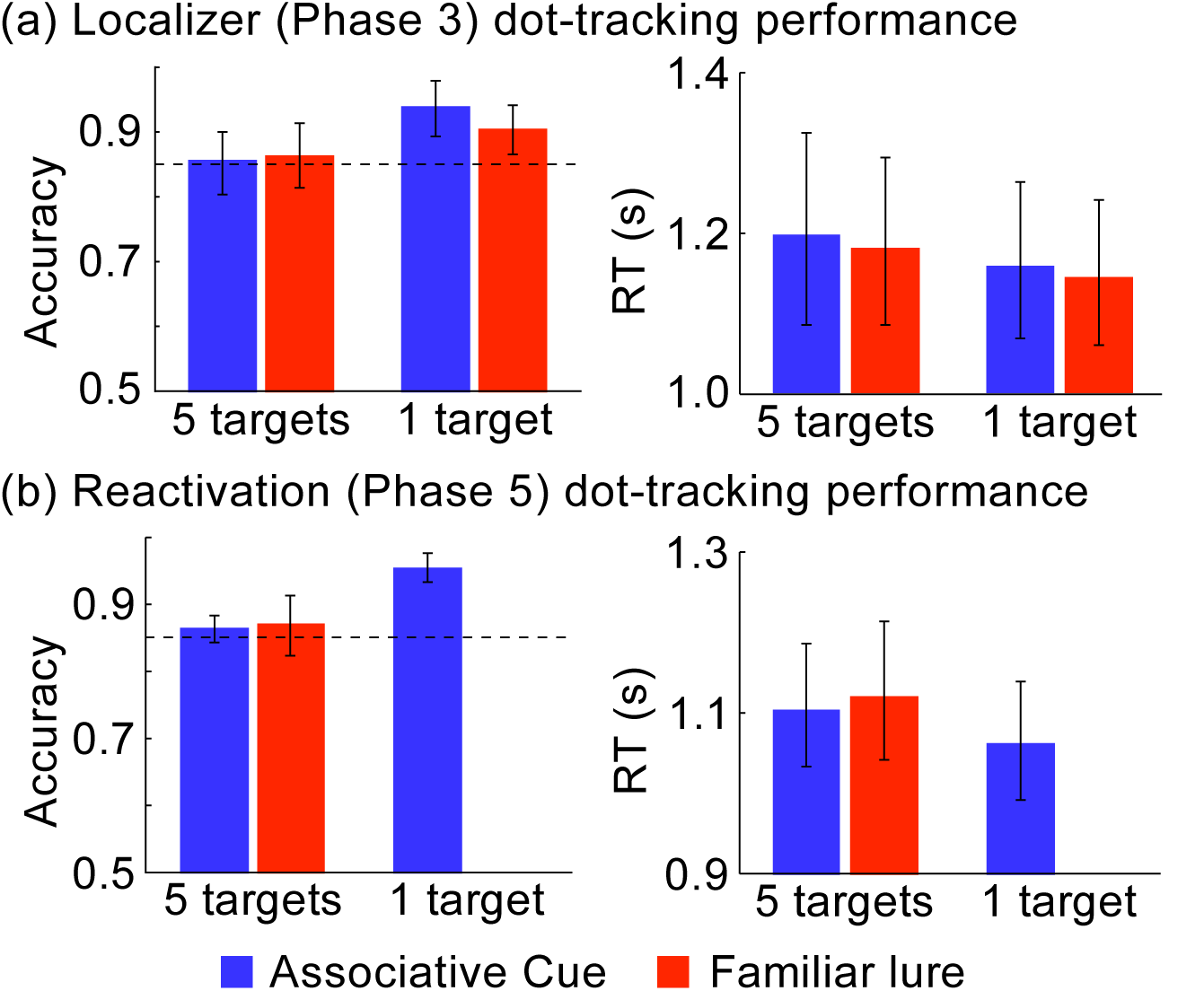
*MOT dot-tracking performance*. Participants were instructed that dot-tracking was their primary task during MOT. During Phase 1, a staircasing algorithm was used to calibrate each participant’s speed of dot movement to a level leading to 85% dot-tracking accuracy during 5 MOT target trials. During both (a) the localizer (Phase 3) and (b) memory reactivation task (Phase 5), dot-tracking performance in 5 MOT target trials remained consistent with this calibrated level. Performance was better for 1 MOT target trials, but no different for associative cue and familiar lure trials. This suggests participants complied with instructions to prioritize dot-tracking, and completed cue visualization using only residual resources, as instructed. Error bars describe 95% CI’s (between-subjects variance; note that comparisons between conditions were performed within-subjects).

### Classifier output as a dependent measure

Having established that we had successfully trained a classifier sensitive to neural evidence of associative recall, we next used this classifier to assess changes in memory reactivation over time. To obtain a temporal “read-out” from a ridge regression classifier corresponding to memory reactivation, we trained a classifier as above using *all* of the data from Phase 3: because brain activity in other phases was of primary theoretical interest, there was no need to create separate training and testing sets within the Phase 3 training data once adequate classifier performance was established. We then used the classifier to independently evaluate each fMRI volume. This yielded, for each timepoint, the amount of evidence in support of the trial being an associative cue trial ("cue evidence") and the amount of evidence in support of the trial being a familiar lure trial ("familiar lure evidence"). We combined these into a single measure by taking the difference between them, and refer to our subtractive measure as "evidence for associative recall". Note that, during classifier training, the target output values for the “associative cue” regression model were perfectly anti-correlated with the target output values for the “familiar lure” regression model (i.e., each trial is either an associative cue trial or a lure trial, never both). Hence, the two regressions might learn mappings whose outputs are perfectly anti-correlated and thus redundant. However, the ridge penalty in ridge regression (which pressures the model to find smaller weights, in addition to minimizing prediction error) exerts an extra effect that – to some extent – decouples the weights of the two classifiers, rendering them non-redundant. As such, taking the difference between outputs has the effect of providing extra information beyond what is obtained from each classifier alone.

The result of our processing was a TR-by-TR (i.e., one 2 s fMRI volume at a time) time-series for each phase, corresponding to a covert measure of associative recall. With this measure established, our next task was to assess the amount of memory reactivation it revealed before MOT-based reactivation, during MOT-based reactivation, and after MOT-based reactivation (Phases 4, 5 and 6, respectively). We accomplished this by extracting the series of values in our classifier output that began just before each memory cue onset, and that ended just before the subsequent event onset. We refer to these time points as TRSTART through TREND. In the Phase 4 and 6 memory tests, *START* corresponded to −1 TR (−2 s) relative to event onset, and *END* corresponded to +5 TRs (10 s) relative to event onset. In Phase 5, *START* corresponded to −1 TR (−2 s) relative to event onset and *END* corresponded to +12 TRs (24 s) relative to event onset. To ensure that we measured evoked, rather than low-frequency state-based signals, we normalized the response to each trial by subtracting the value at trial onset from all TRs within that trial. This baseline was TR0 for Phases 4 and 6, and because extra timepoints were available for Phase 5, it was the average of TR-2 to TR0 in that phase. To reduce the number of comparisons needed for our study, we focused our comparisons on the mean classifier signal om a window of time from 4-8 s for both memory tests, and from 4-18 s for the MOT phase. We started the window at 4 s (instead of 0 s) to account for lag in the hemodynamic response measured with fMRI.

Finally, we organized event responses according to our manipulations. In Phases 4 and 6 (memory testing), we grouped events based on whether the trial belonged to the associative cue (1 MOT target), associative cue (5 MOT targets), or omitted associative cue condition. In Phase 5 (memory reactivation), we grouped events based on whether they belonged to the associative cue (1 MOT target), associative cue (5 MOT targets), or familiar lure (5 MOT targets) condition.

### Significance testing

To provide a random-effects statistical test of condition-level differences, we computed MVPA measures as described above at the single subject level, yielding a different mean memory reactivation timecourse for each condition. Group-level pairwise comparisons of condition means were then conducted using a non-parametric bootstrapping analysis. For each time point, pairwise differences between condition means across participants were calculated. These computations were repeated 10000 times, each time drawing 23 samples with replacement from the group of 23 participants. The standard deviation of differences provided a standard error estimate for each comparison. We divided the overall mean difference by the difference standard error derived from bootstrap resampling to obtain a bootstrap ratio (BSR), which can be treated as an approximate *z* statistic (Efron & Tibshirani, 1986). We set our significance threshold at an absolute value of BSR 1.96 (approximately corresponding to a 95% confidence interval). This same approach was used for the statistical analysis of our behavioral data.

## Results

### Overview

The goal of our experiment was to understand how the Phase 5 difficulty manipulation (1 vs. 5 MOT targets) impacted memory reactivation during associative cue trials, and whether any concurrent impact on later memory (Phase 6) could be ascertained. We also wished to test the usefulness of a novel fMRI classifier trained to measure associative memory and to remain insensitive to the aforementioned difficulty manipulation. We were able to train a classifier that satisfied these properties, and that worked in the context of a moving MOT task. This classifier, as well as participant behavioural responses obtained during the MOT task, indicated that our difficulty manipulation successfully modulated memory reactivation. Evidence from the post-reactivation memory task indicated that the memory representations cued during the MOT task had been weakened, regardless of the level of difficulty.

### Validation of MOT as a scalable distractor task

As discussed, during Phase 1 (staircasing), we used a staircasing method to adjust the speed at which MOT targets moved. We did so in such a way that, when faced with an array of 5 target dots and 5 foil dots, participants could successfully identify a probe dot as either a target or foil 85% of the time. This resulted in a median velocity of 1.43°/s, SD = 0.90°/s, range = [0.52°/s - 3.36°/s]. Each participants’ unique velocity was applied forward throughout their experimental sessions. Our objective for this calibration was to present a similar level of disruption to visualization for all participants. To assess whether our approach was effective, we evaluated participant performance for 5 MOT target trials in the Phase 3 localizer against this 85% accuracy goal. Doing so allowed us to assess whether participants remained engaged throughout the experiment, and did not become substantively better or worse at the task as a result of factors such as ongoing training, fatigue, or the novel fMRI environment. In the Phase 3 localizer task, mean dot-tracking accuracy under 5 MOT target conditions was not significantly different than the staircasing goal of 85% accuracy, BSR = 0.83, P=*n.s.,* range = [75%-100%] (Fig. 2a). In the Phase 5 memory reactivation task, mean dot-tracking accuracy under 5 MOT target conditions was again not significantly different than the staircasing goal of 85% accuracy, BSR = 1.52, *P* = *n.s.,* range = [70%-100%] (Fig. 2b). Although ceiling-level performance in a small subset of participants somewhat complicates interpretation of these values, the results clearly indicate that participants remained engaged throughout the experiment, and suggest that the influence of practice, fatigue and the fMRI environment did not introduce material variation in the executive resources absorbed by the MOT task.

Dot-tracking accuracy data also presented information about the effectiveness of the difficulty manipulation. Performance on 1 MOT target trials was superior to that of 5 MOT target trials, both during the Phase 3 localizer (Fig. 2a), BSR = 3.31, *P* < 0.001, and the Phase 5 memory reactivation (Fig. 2b), BSR = 6.28, *P* < 0.001. Likewise, during Phase 5 memory reactivation, participants were faster to respond (median reaction time; RT) on 1 MOT target trials than 5 MOT target trials, BSR = −2.41, *P* < 0.05, although this pattern was not significant during the localizer, BSR = −1.03, *P* = n.s., which may be attributable to the smaller number of trials contributing to the stability of each participant’s parameter estimates in that phase. These results confirmed that the task was more difficult when it was necessary to track five MOT targets rather than only one.

As a reminder, an important feature of the MOT task was that participants had two competing tasks: dot-tracking, and mental visualization of cued scene associates. For our dot-tracking difficulty manipulation to exert an influence over the amount of memory reactivation experienced by participants, it was important for the dot-tacking task to take priority (i.e., for memory recall to be accomplished using only residual cognitive resources), rather than recall taking priority (i.e., maximizing memory recall, at the expense of dot-tracking performance). Accordingly, we instructed participants to always ensure that dot-tracking remained their top priority during MOT. However, even cooperative participants could have been influenced by automatic processes triggered by a retrieval cue, to the detriment of their dot-tracking performance and our manipulation. To evaluate the extent to which this was an issue, we compared dot-tracking accuracy for cue and familiar lure trials. On cue trials, participants had the opportunity to visualize a scene associate; whereas on familiar lure trials, there was nothing for participants to visualize. In the event that participants did not give full task priority to dot-tracking, then accuracy for cue trials should have been lower than that of lure trials. During the Phase 3 localizer task, we found no such difference in accuracy on 5 MOT target trials, BSR = −0.28, *P* = n.s., nor on 1 MOT target trials, BSR = 1.26, *P* = n.s. Likewise, during 5 MOT target trials in the Phase 5 memory reactivation task, we found no such difference, BSR = −0.30, *P*=n.s. (note: in Phase 5, no lure trials with only 1 MOT target were available for comparison). Dot-tracking RT data (i.e., latency from probe dot presentation to a participant response) also suggested compliance with instructions. During the Phase 3 localizer task, we found no RT differences between cue and lure trials on 5 MOT target trials, BSR = 0.73, *P*=n.s., or 1 MOT target trials, BSR = 0.78, *P*=n.s. During Phase 5 memory reactivation, we also found no differences on 5 MOT target trials, BSR = −0.88, *P*=n.s.

### Validation of classifier measure of memory reactivation

Our classifier performed above chance when tested on left-out portions of the data from the Phase 3 localizer task, *M* = 0.58, 95% CI = [.54 .63], BSR = 3.18, *P <* 0.005. In addition to requiring that our classifier be sensitive to associative recall (i.e., the difference between cues and familiar lures) in the context of a visually dynamic MOT task with variable speeds across participants, an important requirement of our experiment was that, when recall is not taking place, our classifier should be *insensitive* to task difficulty (i.e., number of MOT targets). Although we trained our classifier with these goals in mind, no feature of our design guaranteed that they would be met; it is certainly possible our classifier could track difficulty instead of memory strength. The actual extent to which we were successful in training a classifier that satisfied our goals is an empirical question. Accordingly, we performed a comparison on each fold of cross-validation to establish whether our classifier would distinguish the number of MOT targets on trials when no associative recall was expected to occur (i.e., where memory strength was held constant). In particular, we compared overall classifier evidence for associative recall (i.e., classifier evidence for the word being an associative cue, minus classifier evidence for it being an familiar lure) on familiar lure trials with 5 MOT targets (*M* = −0.04, 95% CI = [−0.10 0.01]) and those with 1 MOT target (*M* = −0.08, 95% CI = [−0.15 −0.02]), and observed no difference, BSR = 1.14, *P* = n.s. (numerically, the difference was in the *opposite* direction from what you would expect if the classifier were confounding increased MOT difficulty with decreased recall). Nonetheless, the classifier was sensitive to MOT task difficulty when associative cues were presented in Phase 5, as we outline in the section below. While we need to be cautious in interpreting null effects, these results support the idea that we had created a classifier that was sensitive to memory strength, but, in the absence of recall, was also insensitive to number of MOT targets.

### Memory reactivation during MOT tasks

We attempted to differentially reactivate memories by varying the number of MOT targets present in a given trial. During Phase 5 (memory reactivation), participants reported lower subjective visualization during associative cue (5 MOT targets) trials than associative cue (1 MOT target) trials, BSR = –2.16, *P* < 0.05 (Fig. 3a), suggesting our manipulation achieved its desired effect. Participants’ subjective responses nonetheless indicated that the MOT task was not so distracting that they were unable to visualize at all, as associative cue (5 MOT targets) trials still had higher-than-null (i.e., a score of 1) visualization, BSR = 24.51, *P* < 0.001. Along these lines, subjective visualization scores for familiar lure (5 MOT target) trials were significantly lower than for associative cue (5 MOT target) trials, BSR = –11.63, *P* < 0.001, and also for associative cue (1 MOT target) trials, BSR = –12.29, *P* < 0.001. These differences indicated that participants’ memories were sufficiently robust for their visualization ratings to discriminate among trials with studied associates (cue trials) and those without (familiar lure trials).

**Fig. 3.**
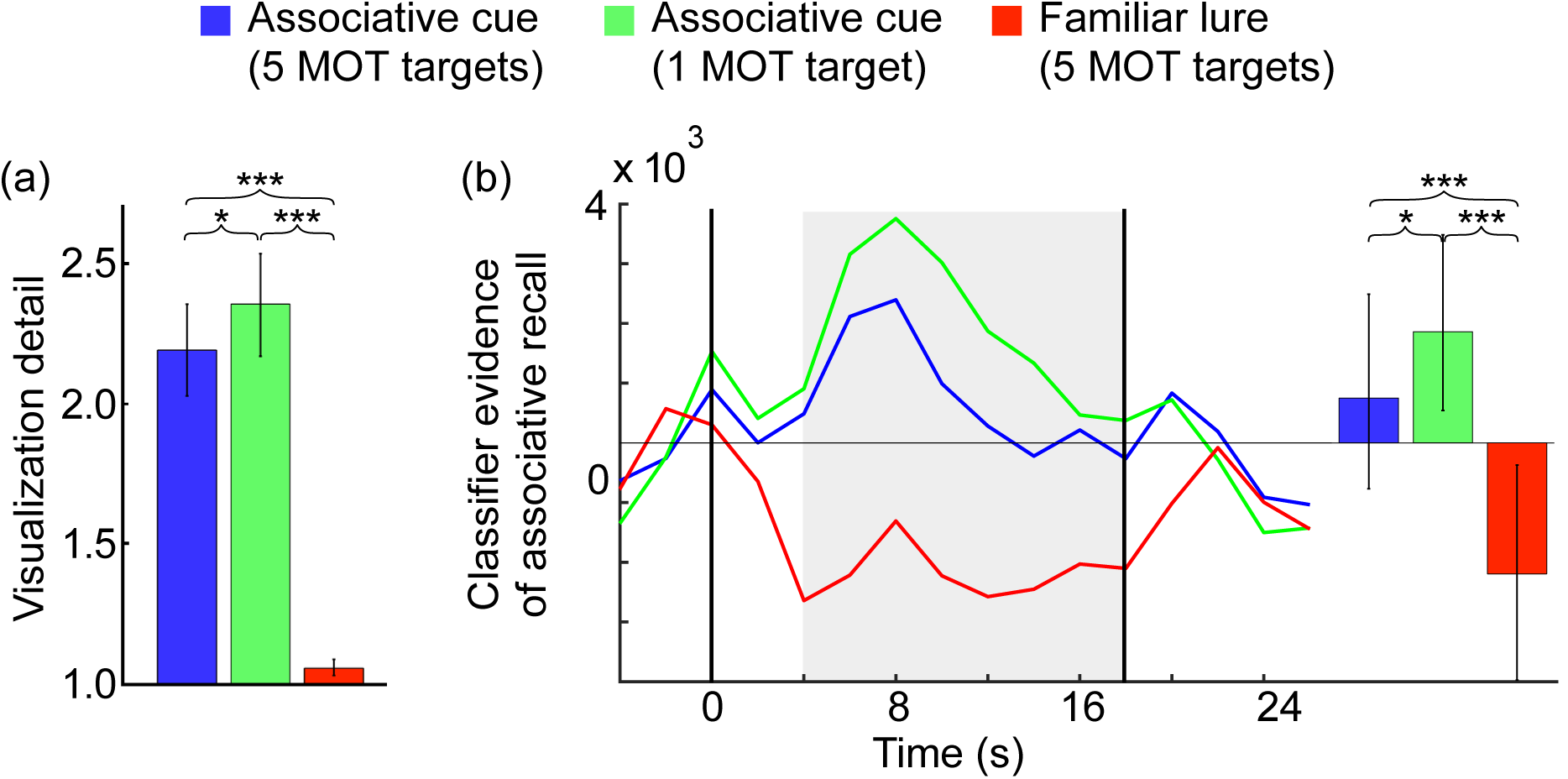
*Evidence of associative recall during the memory reactivation phase*. During the reactivation phase (Phase 5), participants were presented with associative cues (i.e., cues previously associated with scenes) while tracking 5 MOT targets or 1 MOT target, and familiar lures while tracking 5 MOT targets. During tracking, they also reported their subjective visualization of any recalled associate. (a) These ratings were higher for associative cues than familiar lures, regardless of whether 5 or 1 MOT targets were used; but were also significantly higher for associative cue trials with 1 MOT target than ones with 5 MOT targets. (b) When drawing upon classifier evidence of retrieval from this same task, this exact pattern was also observed: classifier evidence from the BOLD-adjusted visualization window (grey; bar plot summary at right) was greater for associative cues than familiar lures, and for 1 MOT target than 5 MOT target associative cue trials. Horizontal lines indicate visualization period onset and offset. Error bars describe 95% CI’s (between-subjects variance; note that comparisons between conditions were performed within-subjects). * indicates BSR > 1.96 (*P* < 0.05); *** indicates BSR > 3.29 (*P* < 0.001).

Participant responses to the four visualization prompts within each MOT trial had low within-trial variance in all types of trials; for 5 MOT target trials, average within-trial variance = 0.19, SD = 0.14; for 1 MOT target trials, average within-trial variance = 0.22, SD = 0.14. It is also worth noting that there was an upwards drift in subjective visualization scores over the course of a trial, which is the opposite pattern to what one would expect if participants were "giving up" on visualization. The mean within-trial slope across the four visualization prompts for associative cue (5 MOT target) trials was 0.03, BSR = 2.49, *P* < 0.05, and the mean within-trial slope for associative cue (1 MOT target) trials was 0.05, BSR = 3.85, *P* < 0.001.

As a heuristic for confirming whether each memory reactivation was partial or full, we compared participants’ subjective evaluation of visualization detail against their original reports during the train-to-criterion task (i.e., after study and prior to reactivation). Because their original scores reflected visualization without distraction, and because these were sampled shortly after study and immediately before correctly identifying the visualized scene in 4AFC, we reasoned that they reflected "full recall". Mean visualization scores during train-to-criterion were 2.64, 95%CI = [2.45 2.86] for the localizer set, and M=2.64, 95%CI = [2.44 2.84] for the recall manipulation set. These scores were higher than were later reported during the Phase 3 localizer task in associative cue (5 MOT target) trials, BSR = 4.19, *P* < 0.001, but not in the associative cue (1 MOT target) trials in that task, BSR = 1.31, *P* = n.s. This confirmed that, during Phase 3, interference from the MOT task induced partial memory reactivation during associative cue (5 MOT target) trials and full reactivation during associative cue (1 MOT target) trials. Likewise, train-to-criterion visualization scores were higher than associative cue (5 MOT targets) trials during the Phase 5 memory reactivation task, BSR = 7.18, *P* < 0.001, (Fig. 3a), although visualization scores were also lower for associative cue (1 MOT target) trials, BSR = 3.62, *P* < 0.001 in that task. Relatively low scores in the associative cue (1 MOT target) condition of Phase 5 are likely attributable to the relatively long study-reactivation interval for the stimulus set in that task (about a day, rather than a few minutes for Phase 3).

Because visualization scores reflect only *subjective* evidence of memory recall, one possible objection to our above findings is that participants’ responses reflected demand characteristics. We therefore sought converging evidence for our manipulation’s effectiveness using an implicit measure of memory reactivation: our trained classifier, which we applied to fMRI data gathered during the MOT phase. Output from the classifier aligned with participants’ subjective reports: greater signal was observed during associative cue (1 MOT target) trials than associative cue (5 MOT target) trials, BSR = 2.05, *P* < 0.05 (Fig. 3b). Also reflecting participants’ reports, classifier output for familiar lure (5 MOT target) trials was significantly lower than associative cue (5 MOT targets) trials, BSR = –3.95, *P* < 0.001, and associative cue (1 MOT target) trials, BSR = – 4.48, *P* < 0.001. All together, the classifier evidence from Phase 5 provided converging support for the idea that partial memory reactivation was modulated by MOT task difficulty. This convergence, in turn, provided a "sanity check" in suggesting the classifier mirrored participants’ own reported memory experiences.

It is worth noting that, because the above analyses average across recall trials, it is possible that evidence of "partial activation" values could arise as an artifact of averaging across "all" and "none" trials. If this were true, we would expect that trial-wise classifier evidence for recall would be bimodally (as opposed to normally) distributed. To test for this, we performed the Shapiro-Wilk parametric hypothesis test of composite normality (which was recently found to be the most powerful normality test in a variety of non-normal situations; Razali & Wah, 2011) on the trial-wise MOT reactivation data from each condition of each participant. The distribution of classifier output across trials did not fit the profile of a bimodal distribution, with the mean of participant *P*-values in the associative cue (5 MOT targets) condition falling well above the cutoff of 0.05 required to assert non-normality, *M* = 0.48, BSR = 7.53, *P* < 0.001. Normality was therefore upheld. This same pattern of high *P-*values was seen in the 1-MOT target condition, *M* = 0.37, BSR = 5.93, *P* < 0.001. Manual inspection of trial-wise histogram data for classifier and cognitive responses further confirmed a normal distribution of reactivation strengths across trials, supporting our interpretation of signal from the MOT phase as reflecting partial memory reactivation.

### Impact of memory reactivation on subsequent recall

To the extent that memories were partially activated during the MOT phase, we hypothesized that this would have a negative impact on subsequent memory performance. To assess this impact, participants were presented with a memory test at the end of the experiment (Phase 6), which investigated memory for associations that had been cued during Phase 5 under associative cue (5 MOT targets), for associative cue (1 MOT target) conditions, and for associations that had not been cued at all during Phase 5. On each trial of the memory test, participants attempted to visualize the scene associate of a cue presented without other distraction, then attempted to select the correct associate from a 4AFC display. Accuracy in the associative cue (5 MOT targets) condition and accuracy in the associative cue (1 MOT target) condition were numerically lower than accuracy for baseline cues (which were omitted from Phase 5 MOT reactivation), but these differences from baseline were not significant: BSR = –1.47, *P* = n.s. for the associative cue (5 MOT targets) condition, and BSR = –0.95, *P* = n.s. for the associative cue (1 MOT target) condition. Likewise, no reliable difference in accuracy was found between associative cue (5 MOT targets) and associative cue (1 MOT target) trials, BSR = –0.29, *P* = n.s (Fig. 4). Implicit measures of memory strength, however, did appear to be impacted. Participants responded more quickly to associative cues left out from the Phase 5 memory reactivation task than associative cues that had been presented with 5 MOT targets, BSR = 2.51, *P* < 0.05, or 1 MOT target, BSR = 2.62, *P* < 0.01. However, there was no difference in response times for associative cue (5 MOT targets) or associative cue (1 MOT target) trials, BSR = 0.38, *P* = n.s. Classifier evidence showed a similar pattern (Fig. 4d): reduced classifier signal was observed in association with associative cue (5 MOT targets) trials relative to ones left out from Phase 5 memory reactivation, BSR = –2.58, *P* < 0.01. No such difference was found for associative cue (1 MOT target) trials, BSR = –1.47, *P* = n.s., and there was also no significant

**Fig. 4.**
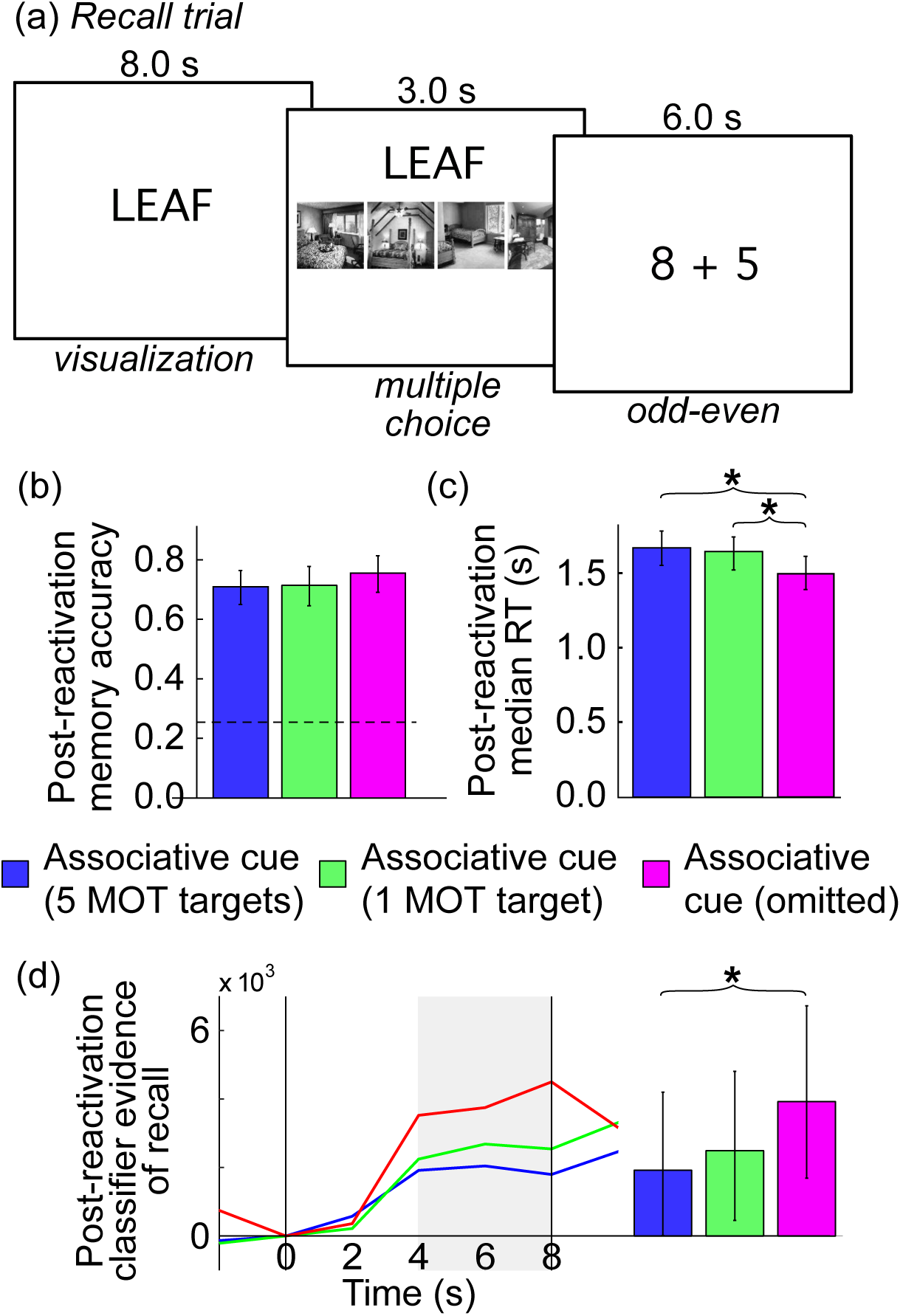
*Impact of memory reactivation on later memory.* (a) Both before and after the memory reactivation phase, participants completed a memory test in which they first visualized the scene associate of memory cues, then completed a multiple-choice question, followed by a mathematical distractor task. (b) Memory accuracy was numerically, but not significantly lower for cues that had been presented during the MOT reactivation phase (dashed line reveals chance performance level). (c) Response times were significantly slower for cues that had been presented during the MOT reactivation phase. (d) Classifier evidence from the BOLD-adjusted visualization window (grey; bar plot summary at right) was lower for cues that had been presented alongside 5 MOT targets during the MOT reactivation phase. Horizontal lines indicate visualization period onset and offset. Error bars describe 95% CI’s (between-subjects variance; note that comparisons between conditions were performed within-subjects). * indicates BSR > 1.96. difference in classifier output for associative cue (5 MOT targets) and associative cue (1 MOT target) trials, BSR = –0.72, *P* = n.s.

## Discussion

In the current study, we sought to establish a parametrically scalable procedure for reactivating memories. Our first contribution was to implement a procedure that, according to both classifier evidence as well as participant subjective reports, was successful both at partially activating memories and modulating the particular amount of partial activation that took place. As predicted, this procedure led to evidence of memory weakening in a post-reactivation memory test, although for more definitive tests of the non-monotonic plasticity hypothesis, it will be necessary to select parameters that broaden the range of observed partial reactivation values. Our second contribution was to train an "associative recall" classifier able to distinguish cues with associates from familiar lures, while remaining insensitive to irrelevant factors (such as MOT difficulty).

### Parametrically scalable memory reactivation

As we have argued, the experimental procedures used to study memory weakening typically incorporate binary manipulations (e.g., retrieval practice; and think/no-think paradigms). Across many studies, these manipulations have been shown to lead to weakening (Murayama et al., 2014); according to the non-monotonic plasticity hypothesis, this is because they induce intermediate levels of memory reactivation (e.g., Newman & Norman, 2010; Lewis-Peacock & Norman, 2014). However, "partial memory reactivation" is not a discrete state; rather, memory reactivation and its downstream effects fall on a continuous dimension (e.g., Johnson, McDuff, Rugg, & Norman, 2009; see also Detre et al., 2013, for evidence and discussion). Here, we have shown an MOT difficulty-based manipulation to be effective at influencing memory reactivation in a graded manner. In particular, altering the number of target dots to be tracked in an MOT task while participants concurrently performed mental visualization allowed us to: 1) reduce memory reactivation below baseline levels on a behavioural index; and 2) modulate fMRI classifier evidence of memory reactivation.

In the current study, we chose to manipulate MOT difficulty by modulating the number of target dots that participants needed to track during dot-tracking. This had the advantage that perceptual features were nearly identical across difficulty conditions, with the only difference between "easy" and "hard" trials being the number of dots painted as MOT targets *prior to* the onset of the trial. A limitation of manipulating the number of MOT targets is that it can only be manipulated in discrete steps (adding or removing an MOT target) – in our experiment, one MOT target still imposed sufficient processing load to induce less-than-full memory reactivation. As such, future work might benefit from other, more fine-grained ways of manipulating difficulty. Notably, prior work has found that it is principally the amount of time that tracked MOT targets spend in close proximity to lures that consumes executive resources (Franconeri, Jonathan, & Scimeca, 2010; Franconeri, Lin, Enns, Pylyshyn, & Fisher, 2008). Changes such as increasing dot speed, growing the size of dots relative to the area they can move on the screen, increasing dot clustering behaviour, or altering other parameters that increase the frequency of dot collisions are therefore expected to have similar resource-depleting effects to our own difficulty manipulation of increasing the number of MOT targets. Accordingly, these parameters should have similar effects on mental visualization if used in conjunction with a reactivation task. Modifying these parameters to influence memory reactivation may be advantageous in that they lie on a truly continuous distribution (unlike manipulating the number of dots that are MOT targets) and thus can be adjusted to induce a broader range of task difficulty values.

It should be acknowledged that the multi-faceted nature of the task made it difficult to explain and perform, but with coaching, practice, and calibration of dot velocity to the individual ability, participants were able to master it. In particular, they showed high accuracy on the MOT dot classification task, which requires vigilance throughout the entire trial period, alongside stable visualization reports during MOT trials. These reports showed a slight upwards bias (i.e., more visualization over time). Together, these observations suggest that participants remained engaged in and could adequately perform both aspects of the task.

### Effects of partial reactivation on memory

Our study joins a growing number of experiments that have illustrated a link between classifier evidence of partial memory reactivation and weaker overall subsequent memory (e.g., Detre et al., 2013; Lewis-Peacock & Norman, 2014; Poppenk & Norman, 2014). Memory cues that were exposed during MOT – whether participants were under instructions to track 5 MOT targets or just 1 MOT target – were shown, on average, to be partially activated. Relative to other memory cues not presented during that phase, memory for the partially activated items was found to be weakened in a post-reactivation memory test, as revealed by both slower response times and lower classifier output than in the pre-reactivation memory test. Numerically, weakening (i.e., a reduction in memory strength relative to the omit / not-reactivated condition) was consistently greatest across our dependent measures (accuracy, reaction time, and classifier output) for associative cue (5 MOT targets) trials, which was also the only condition to show significant classifier evidence of weakening. However, none of these variables revealed significant differences when associative cue (5 MOT target) and associative cue (1 MOT target) trials were compared directly; and associative cue (1 MOT target) trials *did* show significantly slower reaction times than omitted items. This pattern likely reflects the fact that during the MOT task, participants’ subjective ratings indicated that reactivation was less than "full" even for associative cue (1 MOT target) trials; and that although this partial reactivation pushed items somewhat out of the reactivation range associated with weakening, some weakening nonetheless took place. The pattern also limits the strength of the argument that can be made about the impact of partial reactivation on forgetting, as when interpreted in isolation, it leaves open the logical possibility that reactivation *in general* causes weakening. The present study can best be viewed as a “proof of concept” that memory reactivation strength can be parametrically manipulated using MOT, leading to memory weakening. In future work, we will parametrically vary reactivation across a wider range of values, with the goal of fully reconstructing the U-shaped curve predicted by the non-monotonic plasticity hypothesis.

### Associative recall classifier

Training a classifier capable of measuring memory reactivation in the context of our new procedure was challenging, as conventional, visual category-based classifiers appeared to attribute the moving MOT dot fields as similar to a particular visual category ("scenes"). We solved this issue by using a procedure similar to that of Rissman et al. (2010): training our classifier on the basis of memory operations (associative recall using cue words vs. recognition of familiar lures) rather than the more typical approach of using distinctive visual categories (e.g., Spiridon & Kanwisher, 2002). By supplying the classifier with trials that varied in difficulty within the same condition, we ensured that training on difficulty-linked features would yield low classifier accuracy, and reduced the probability that classifier output would be sensitive to these features. We found that, when MOT task difficulty was held constant, this classifier was able to deliver above-chance performance in the challenging cognitive environment of dot-tracking in an MOT task. The trained classifier also met the important requirement of being insensitive to task difficulty when memory cues were not present. This pattern indicates that differences in classifier evidence evoked by associative cues with 1 vs. 5 MOT targets reflected different levels of memory retrieval strength, rather than task difficulty per se.

### Applications

We anticipate that there will be many uses for paradigms like this one that provide greater control over levels of memory reactivation. We wish to highlight two important applications of interest to our own laboratories. First, as noted earlier, experiments aimed at charting the "link function" between memory reactivation and subsequent memory strength (e.g., Detre et al., 2013) have relied on naturally-occurring variability in memory activation strength. A shortcoming of this approach is that mapping out the full U-shaped curves requires observations at a wide range of recall strength levels and there is no guarantee that enough observations will be obtained at these levels (especially at the high and low extremes). By exposing participants to a range of MOT difficulty parameters that yield lower and higher memory reactivation, it may be possible (in future work) to use the paradigm described here to populate the tails, and therefore sample from a more uniform memory reactivation distribution.

Along these lines, another important affordance of a parametrically scalable reactivation protocol is the possibility of adapting it towards closed loop experimentation, adjusting difficulty as each trial unfolds in an attempt to generate memory reactivation at particular levels. A classifier in an fMRI environment configured to deliver a live read-out (e.g., deBettencourt, Cohen, Lee, Norman, & Turk-Browne, 2015) could, in the context of the current procedure, provide information about the amount of memory reactivation triggered by the current memory cue at the current level of MOT difficulty, accounting for variation injected by fluctuations in the association’s strength and the participant’s attention. This information, in turn, could be used to modulate difficulty levels such memory reactivation could be readjusted towards a goal level. This introduces the possibility of a causal test of the non-monotonic plasticity hypothesis: experimenters could induce partial memory reactivation at specific sections of the non-monotonic plasticity curve, probing for predicted impacts on subsequent memory.

Eventually, therapeutically applied versions of closed-loop procedures could be used to steer *all* memories into the portion of the non-monotonic plasticity curve most associated with weakening, with the goal of attenuating the traumatic associates of powerful memory cues. Most phases of our design could be eliminated this context, since patients would not need to learn new associations – presumably, the traumatic associations would precede therapy. Only MOT difficulty calibration (Phase 1), localizer training (Phase 3), and memory reactivation (Phase 5) would be required. As these steps could easily be completed in two short sessions, we believe our technique to be viable as a prospective therapeutic approach.

### Conclusions

In summary, we have illustrated a "proof of concept" application of an MOT-based procedure for parametrically modulating memory reactivation. Behavioural and classifier measures of reactivation both confirmed that modulating MOT difficulty influenced the degree of memory reactivation. In turn, this partial memory reactivation appeared to lead to subsequent memory weakening. This procedure is intended to make possible new, focused investigations into human learning that exert greater experimental control over memory reactivation to conduct, for example, causal tests of the non-monotonic plasticity hypothesis. Our procedure also may pave the way for closed-loop clinical procedures that are based on principles of partial memory reactivation.

